# PhOxi-seq detects enzyme-dependent m^2^G in multiple RNA types

**DOI:** 10.1101/2024.08.15.608094

**Authors:** Marie Klimontova, Kimberley Chung Kim Chung, Han Zhang, Tony Kouzarides, Andrew J Bannister, Ryan Hili

## Abstract

In recent years, RNA-modifying enzymes have gained significant attention due to their impact on critical RNA-based processes, and consequently human pathology. However, identifying sites of modifications throughout the transcriptome remains challenging largely due to the lack of accurate and sensitive detection technologies. Recently, we described PhOxi-seq as a method capable of confirming known sites of m^2^G within abundant classes of RNA, namely purified rRNA and purified tRNA. Here, we further explore the selectivity of PhOxi-seq and describe an optimised PhOxi-seq workflow, coupled to a novel bioinformatic pipeline, that is capable of detecting enzyme-dependent m^2^G sites throughout the transcriptome, including low abundant mRNAs. In this way, we generated the first database of high confidence sites of THUMPD3-dependent m^2^G in multiple RNA classes within a human cancer cell line and further identify non-THUMPD3 controlled sites throughout the transcriptome.

---

From their first identification in the early 1950’s to present, over 150 post-transcriptional RNA modifications have been identified across all life.^1^ These chemical modifications intricately regulate diverse functions of RNA molecules and govern root cellular processes ranging from splicing, translation, and degradation.^2^ Not surprisingly, mounting evidence underscores their pivotal roles in the etiology of human disease, particularly cancer.^3^ Sequencing technologies that can map these RNA modifications transcriptome-wide at single-nucleotide resolution have been instrumental in guiding functional studies.^4^ Such methods have been critical to the developing picture of several prevalent RNA modifications, such as *N*^6^-methyladenosine (m^6^A), and its role in RNA biology^5^ and human diseases.^6^ However, a general lack of accurate and sensitive methods to map and quantify most RNA modifications has significantly hampered the development of this burgeoning field of research. Indeed, most existing techniques rely on some form of modification-specific immunoprecipitation of RNA coupled to next-generation sequencing. Unfortunately, these methods necessarily depend upon the availability of specific antibodies, which are limited or simply not available for many RNA modifications. Consequently, there is a pressing need for novel antibody-independent approaches to expedite investigations into the functions of RNA modifications.

*N*^2^-Methylguanosine (m^2^G) has been identified in various RNA molecules across all three domains of life: Archaea, Bacteria, and Eukarya. While most often associated with tRNA and rRNA, m^2^G was recently detected by mass spectrometry analysis of mRNA isolated from *S. cerevisiae*, suggesting that it may have regulatory functions at the transcriptome level, and could be present in other species.^7^ Recently, three m^2^G-catalyzing RNA methyltransferases – THUMPD2, THUMPD3, and TRMT11 – have been characterized.^8-10^ The dysregulation of these methyltransferases have been associated with various human diseases. THUMPD2 is known to install a critical m^2^G modification in *U6* snRNA, the catalytic centre of the spliceosome, necessary for efficient splicing activity; its dysregulation is linked to age-related macular degeneration and retinal function.^10^ Upregulated THUMPD3 has been shown to promote lung cancer cell proliferation and migration by modulating the extracellular matrix through mechanisms such as alternative splicing of protein transcripts.^11^ Lastly, TRMT11 fusions have been associated with aggressive and recurring prostate cancers.^12^ Further functional studies are required to determine the role of m^2^G in human health; however, limited tools are available to experimentally study m^2^G. Early transcriptome maps of other post-transcriptional modifications, such as m^6^A, relied on established antibodies, well-characterised demethylases, and a relatively high natural abundance of the modification.^13^ On the contrary, m^2^G lacks these essential tools making its study particularly challenging.

We previously described photooxidative sequencing (PhOxi-seq), which enables the single-nucleotide resolution sequencing of m^2^G by harnessing the chemoselectivity of photoredox chemistry to selectively photooxidize m^2^G in the presence of guanosine.^14^ The method relies upon the lower oxidation potential of methylated guanine over that of canonical guanine. In the presence of blue light, selectfluor, and riboflavin as a photocatalyst, m^2^G is mutated into m^2^-Iz and m^2^-OG, which pair with G and A, respectively (Figure 1). PhOxi-seq was shown to readily identify m^2^G sites in model synthetic RNAs, as well as known sites in highly abundant RNAs, such as purified yeast tRNAs and purified *E. coli* rRNA. Given the limited understanding of the biological role of m^2^G and the potential of m^2^G methyltransferases as therapeutic targets, we sought to adapt PhOxi-seq to establish the first putative m^2^G map of the transcriptome to facilitate their functional studies.

**Figure 1.**
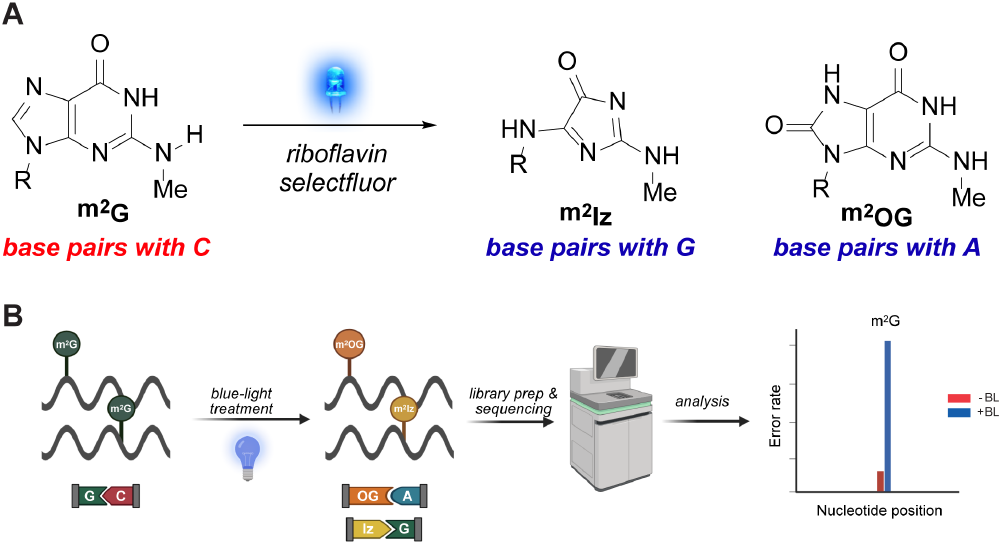
A) Photooxidation reaction of N^2^-Methylguanosine into N^2^-imidazolone (m^2^Iz) and N^2^-oxoguanine (m^2^OG), which is the chemical basis for photooxidative sequencing (PhOxi-seq). B) PhOxi-seq workflow to detect m^2^G in RNA.

## RESULTS AND DISCUSSION

During the development and validation of novel sequencing methods for RNA modifications, false positives are reduced by using antibodies to enrich the modification, or demethylases are employed to remove the modification to confirm the signal. Lacking these important tools, we took inspiration from m^6^A sequencing methods that knocked down individual methyltransferases to establish which m^6^A sites in the human transcriptome were under the control of a specific methyltransferase.^15-19^ To this end, we investigated the multiple known m^2^G RNA methyltransferases responsible for installing m^2^G within RNA molecules to facilitate m^2^G identification.

We first considered whether PhOxi-seq could be used to detect specific enzyme-dependent sites of m^2^G within tRNAs. We previously used PhOxi-seq to identify m^2^G within tRNAs isolated from yeast.^14^ Other works using mass spectrometry-coupled methods later showed that human THUMPD3 installs m^2^G at position 6 within specific tRNAs, including isodecoders *tRNA-Gly-GCC-2* and *tRNA-Gly-CCC-2*. Importantly, m^2^G6 within these isodecoders was shown to be significantly decreased by loss of THUMPD3.^8^ We first asked whether PhOxi-seq could detect m^2^G6 within these particular tRNA isodecoders. To ensure accurate analysis of m^2^G6 within the specific tRNA isodecoders, we designed specific primers for the respective decoders (Figure S1A and S1B). YAMAT-seq^20^ confirmed accurate targeting of the relevant isodecoders (Figure S1C). Our data clearly show that PhOxi-seq detects m^2^G6 within *tRNA-Gly-GCC-2* (Figure 2A, WT) and *tRNA-Gly-CCC-2* (Figure S1D, WT). Next, we asked whether PhOxi-seq could identify THUMPD3-dependent sites of m^2^G within the same tRNA iso-decoders. We compared the sequencing error frequency following blue light treatment (+BL) of RNA samples harvested from WT cells and THUMPD3-depleted (D3kd) cells (Figure 2A and S1D). The data reveal significantly reduced sequencing error frequencies at G6 of *tRNA-Gly-GCC-2* and *tRNA-Gly-CCC-2* in tRNA samples isolated from THUMPD3-depleted cells, indicating that THUMPD3 catalyzes a significant proportion of m^2^G at this site. Remaining m^2^G signal may be the result of compensatory methylation by another guanine methylase or may be ascribed to the creation of a hypomorphic THUMPD3 allele, which has been frequently observed in METTL3 CRISPR knockouts during m^6^A studies.^21^ As expected with guanosine oxidation pathways, we observed T and C substitutions as significant nucleotide sequencing errors at the known m^2^G position (Figure 2B and S1E). Interestingly, in addition to the expected substitutions, another error type, G skipping, was prevalent. This identified a novel type of sequencing error that can be used in future analysis to detect genome-wide m^2^G distribution. Taken together, these findings highlight PhOxi-seq as a technique capable of detecting enzyme-dependent sites of m^2^G within tRNA. Furthermore, the unexpected error rate profile may prove helpful in identifying other potential THUMPD3-dependent sites of m^2^G. Based on these findings, we decided to explore whether PhOxi-seq could be used to detect m^2^G sites within the wider transcriptome.

**Figure 2.**
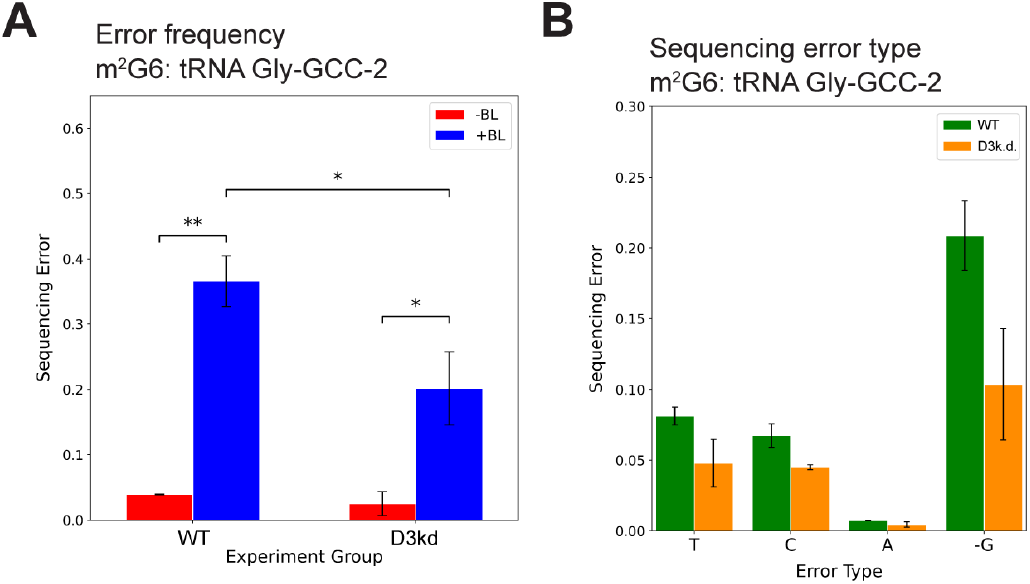
THUMPD3-depletion in A549 lung cancer cells results in reduced m^2^G6 in tRNA, detected via PhOxi-seq. A) Plot representing sequencing error at G6 of tRNA-Gly-GCC-2 with (+BL; blue) and without (-BL; red) blue-light treatment. Two experimental groups represent control (WT) and THUMPD3-depleted cell lines (D3kd). Error bars represent the mean ± SD of 3 independent replicates (n=3). Welch’s t-test was used to assess the significance of changes in error rates at G6 on tRNA-Gly-GCC-2. ns - P > 0.05, * - P ≤ 0.05, ** - P ≤ 0.01. B) Sequencing error data represented by the blue bars (+BL samples) were deconvoluted into individual nucleotides. Respective proportion of reads corresponding to main types of misincorporation – G-T substitution, G-C substitution, G-A substitution, and single base deletion were plotted. Green and orange bars represent WT and D3k.d. data, respectively. Error bars represent the mean ± SD of 3 independent replicates (n=3).

Our initial findings validate PhOxi-seq as an effective method of identifying enzyme-dependent sites of m^2^G within specific tRNAs. However, its application and relevance to mRNA and other non-coding RNAs has not been explored. Based on our prior success with tRNA and rRNA, we reasoned that blue light-induced photooxidation of m^2^G is independent of RNA type. Therefore, to identify sites of m^2^G within other types of RNA, we subjected polyA+ enriched RNA to the same photooxidation condition and subjected the resulting RNA libraries to high throughput sequencing. However, as PhOxi-seq has not previously been tested on this class of RNA, we had to generate a unique bioinformatic pipeline (Figure 2A and S2).

As a relatively crude overview of the data to determine if THUMPD3 depletion affected error rates in a subset of reads, we calculated density plots of variant allele frequency (VAF) data distribution for each sample (Figure 3B). VAF data encompasses single-nucleotide polymorphisms (SNPs) and other single-nucleotide variations from the reference genome (*i*.*e*., sequencing error rates). The plots clearly show reduction of variant sites with VAF values between 0.1 and 0.5 upon THUMPD3 depletion. These data are consistent with loss of blue-light induced changes at sites of m^2^G when THUMPD3 is targeted. At the end of the bioinformatic pipeline, prior to the target identification, we analysed whether low sequencing depth leads to more uncertainty of the quantification of low abundance genes, as might be expected from results in both bulk and single cell sequencing technologies. We therefore calculated the average sequencing depth of all SNPs across different samples. These were then sorted, in ascending order, by coverage and we employed a non-overlapping sliding window approach to group the sorted sites into ‘windows’. We then calculated Spearman’s rank correlation coefficient and plotted the data as shown in Figure 3C. This shows that correlations increase with higher sequencing depth, providing a rational to filter out low read depth regions (indicated by dotted line, Figure 3C; read depth ≥ 50). This cut-off was included in our downstream analysis of the pipeline data (Figure 3D). The rationale for these cut-off limits is described in the supporting information; however, it is worth noting here that limiting VAF in samples without BL treatment to ≤ 5% eliminated possible signals from m^2^_2_G, which is sensitive to PhOxi-seq, albeit with a large observed decrease in error.^14^

**Figure 3.**
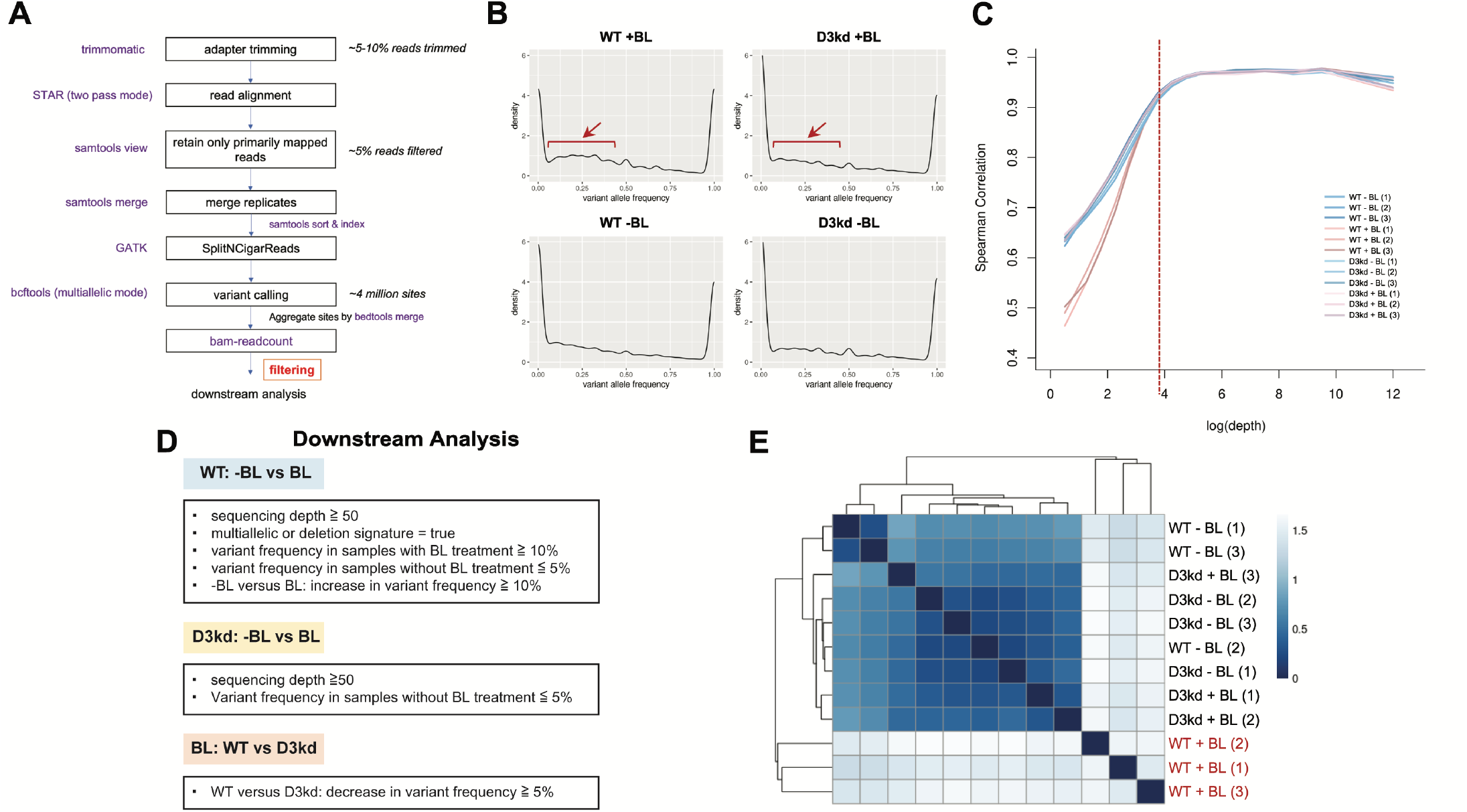
Overview of quality control (QC) measures of outputs from the PhOxi-seq pipeline on the whole transcriptome. (A) Overview of PhOxi-seq pipeline on the whole transcriptome. Main steps of the pipeline were presented in flowchart, involving steps of read processing, alignment, variant calling, and downstream analysis. Softwares involved at each step were labelled on the left and coloured purple, percentage of reads processed at each step and number of variant sites were labelled on the right. The conditions of filtering step coloured in red could be found in Figure 3D. The detailed description, rationale and quality control steps of the pipeline could be found in Supporting Information. (B) Representative density plots from one biological replicate illustrating overall distribution of variant allele frequency (VAF) data of variant sites of representative replicate from each experiment group (control cell line (WT) with and without BL (±BL) treatment, THUMPD3 depleted cell line (D3kd) ±BL treatment). The y-axis of the density curve is density and the area under the curve gives the percentage. The x-axis ranges from 0-1, representing the VAF data, which here indicates the proportion of reads that do not match the reference genome at each variant site. The consistent pattern in the plots with two peaks correspond to specific sets of data: one peak with VAF values (x-axis) approaching 1 represent mostly SNPs; one with VAF values approaching 0 could be mostly noises. The VAF data in this plot covers variant sites from the variant calling step that show non-zero depth in all replicates (n=1192693). (C) Line plot of average Spearman’s rank correlation coefficient (y-axis) of VAF data for each replicate with the rest of the replicates of increasing windows of average sequencing coverage (x-axis). The y-axis ranges from 0.4 – 1. The closer the coefficient values to 1, the stronger the correlation, and vice versa. The average sequencing depth of all ‘SNP’ sites across different samples were calculated. These sites are then sorted, in ascending order, by coverage. A non-overlapping sliding window approach is used to group the sorted sites where sites with similar coverage in a specific range are grouped into ‘windows’. For each window (e.g. log(depth): 2-3, 3-4, log here represents natural log), the Spearman’s rank correlation of the VAF data of the sites from the sample of interest and all other samples is calculated and averaged, and plotted. A vertical dotted line indicates the depth threshold of 50 at x-axis = 3.912 (log(50)). (D) Table illustrating filtering conditions for potential m^2^G modified sites during downstream analysis. The conditions include threshold on sequencing coverage, levels of sequencing error changes in different groups and presence of multiallelic signature. Further details are noted in the supporting information. (E) Heatmap of the sample-to-sample distances based on Euclidean distance measure for sequencing error rates of top potential m^2^G sites (n=62) of all samples. A hierarchical clustering was also performed based on the sample distances and presented in the plot.

We next assessed the relationship between VAF of each sample. Our analysis revealed that blue light-treated RNA from WT cells exhibited the greatest differences from all other samples (Figure S2B). When only the top high confidence targets were considered, this difference was even more pronounced (Figure 3E), which is consistent with the expected higher sequencing error frequency in samples with intact m^2^G prior to THUMPD3 depletion. Overall, our findings suggest that PhOxi-treatment induces sequencing error rates that are largely resolved upon THUMPD3 depletion, underscoring the utility and effectiveness of PhOxi-seq in identifying enzyme-dependent m^2^G.

Finally, we leveraged PhOxi-seq and our novel bioinformatic pipeline to identify sites of m^2^G within a human polyA+ enriched transcriptome isolated from A549 lung cancer cells. Analysis of our tRNA data revealed that each m^2^G site treated with photooxidation mutates to any base (although with different frequencies, Figure 2B) or gets deleted. Applying these characteristics where any change or deletion of G is scored within our pipeline, we identified 381 potential THUMPD3-dependent sites of m^2^G within 298 distinct polyA+ enriched RNAs (Table S1).

Since multiple base substitutions and deletion signatures were observed at m^2^G6 in tRNAs, we applied to the pipeline a requirement to further filter our hits for at least two photooxidation induced mutational changes at the sites identified above (Figure 2B). This revealed 62 high confidence hits (Figure 4A and Table S2). These m^2^G sites were distributed across various mRNA regions, including exons (15%), introns (61%), 3’ UTR (11%) and 5’ UTR (2%) (Figure S5). Interestingly, m^2^G sites were enriched within intronic regions. This aligns well with the previously reported role of THUMPD3 in the regulation of RNA splicing.^11^ Consistent with previous observations that elevated THUMPD3 is associated with cancer phenotypes,^11^ several putative m^2^G sites were located on oncogenic transcripts. A notable example is integrin β4 subunit (ITGB4), which has been associated with several cancer types, including lung carcinoma,^22^ and showed a marked response in PhOxi-seq for a putative m^2^G site in an exon location (Figure 4B). This signal was significantly deceased for THUMPD3-depleted samples (Figure 4C). Several putative m^2^G sites appeared to be highly dependent on THUMPD3, such as that observed in the intron of the E3 ligase Thyroid hormone receptor interactor 12 (TRIP12), which experienced complete loss of photooxidation-mediated mutation in THUMPD3-depleted cells (Table S2). Interestingly, TRIP12 is often mutated in lung adenocarcinoma patients,^23^ and is known to regulate the extracellular matrix.^24^ Notably, our analysis also uncovered m^2^G sites within a well-characterised non-coding RNA (*7SK*; Tables S1 and S2; Figure S3C).

**Figure 4.**
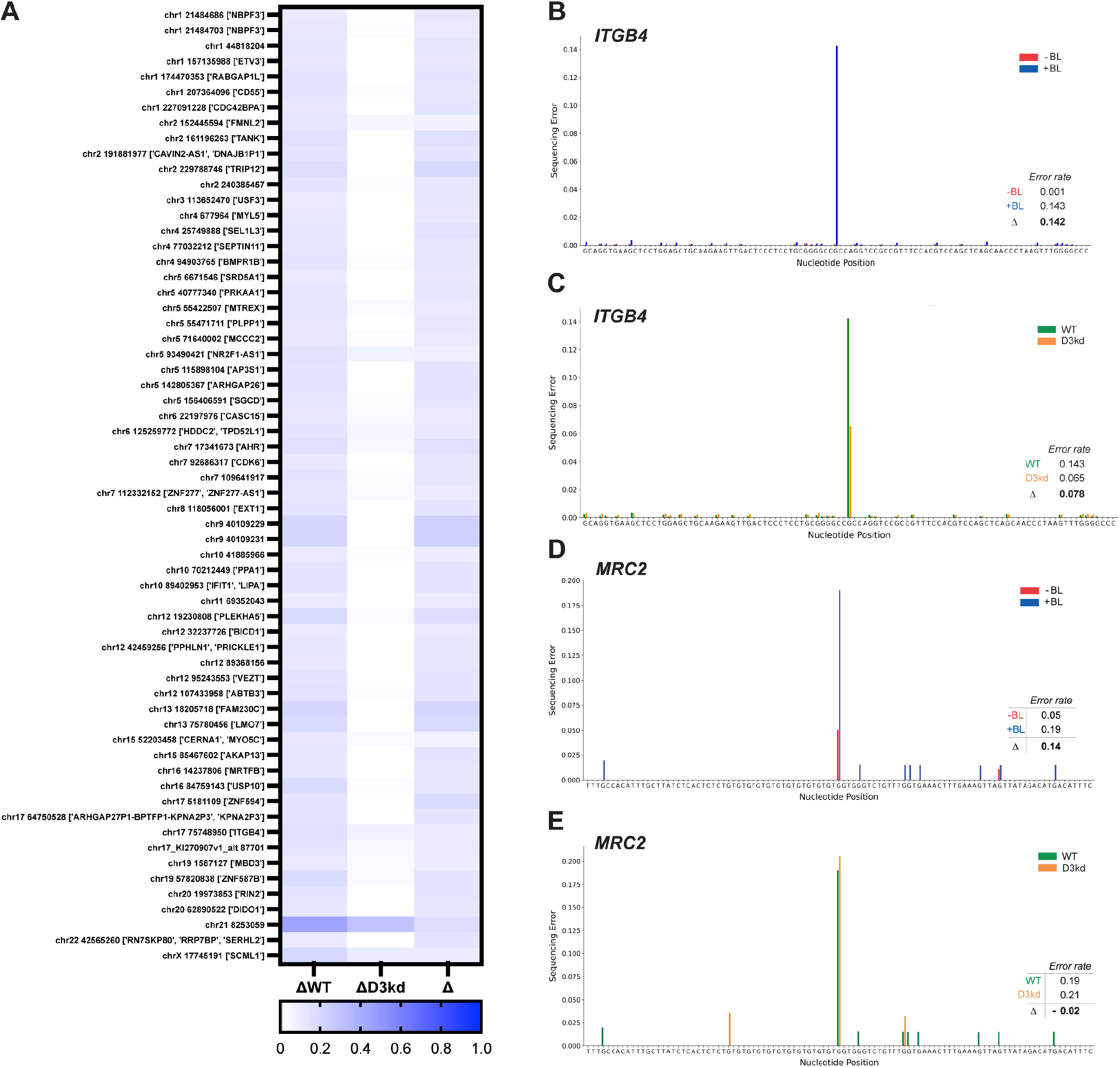
Effect of PhOxi-seq treatment on total sequencing errors in the genomic regions of top potential THUMPD3 dependent and independent m^2^G sites. A) heatmap of high-confidence THUMPD3-dependent m^2^G sites. Transcript and location are plotted with difference in PhOxi-seq response in wild type A549 lung cancer cells (ΔWT), difference in PhOxi-seq response in THUMPD3 knockdown A549 lung cancer cells (ΔD3kd), and the difference between the ΔWT and ΔD3kd experiments (Δ). B) Plot representing guanosine error in THUMPD3-dependent site, ITGB4 for samples with (+BL, depicted in blue) and without (-BL, depicted in red) blue-light treatment. C) Bar plot of guanosine errors in ITGB4 for the photo-oxidised samples (+BL) in WT and THUMPD3-depleted (ΔD3kd) cells. D) Plot of guanosine error in THUMPD3-independent site, MRC2, for samples with (+BL) and without (-BL) blue-light treatment. E) Guanosine error rates in MRC2 for the photo-oxidised samples (+BL) in WT and THUMPD3-depleted (ΔD3kd) cells. Green bars indicate WT cell lines and orange bars indicate THUMPD3 depleted cell line D3kd). The sequencing error is illustrated in percentage, where 0 is 0% and 1 represents 100%. Insert in graph: tables indicating error rates at given m^2^G sites.

Beyond the putative THUMPD3-controlled m^2^G sites we identified above, we identified a total of 2978 or 362 blue-light sensitive G sites (lower and higher confidence hits, respectively) which are based on only WT samples (Tables S3 and S4). This list comprises both THUMPD3-dependent and THUMPD3-independent sites. Additional sites that were unaffected by THUMPD3 depletion suggests the presence of a THUMPD3-independent modification at these sites. For example, a potential THUMPD3-independent m^2^G site is located in the mannose receptor C-type 2 transcript (MRC2), which does not resolve photooxidation-induced error in THUMPD3-depleted samples (Figures 4D and 4E). Such sites may be under the control of another m^2^G methylases, such as THUMPD2 or TRMT11.

## CONCLUSION

Herein, we have described an optimised PhOxi-seq method that we used to provide the first putative map of THUMPD3-dependent m^2^G in various RNA species. This involved optimising the technique to reliably confirm known sites of THUMPD3-dependent m^2^G in tRNA and then considerably extended the scope of PhOxi-seq to detect THUMPD3-dependent m^2^G sites in other RNA species.

Given the lack of tools to study m^2^G, we sought to establish high confidence m^2^G sites by knocking down THUMPD3, a known methyltransferase for m^2^G installation in RNA. Our analysis of THUMPD3-dependent sites of m^2^G within tRNA indicated that PhOxi-seq could detect such sites with a good signal-to-noise ratio. Furthermore, the analysis of the tRNA data involved the creation of an advanced bioinformatic pipeline to identify enzyme-dependent sites of modification. This was fundamentally important before other types of RNA could be analysed.

At present, the identification of specific sites of a given modification within less abundant RNAs remains a significant challenge within the RNA modification community. Consequently, we applied PhOxi-seq coupled with our novel bioinformatic pipeline to detect THUMPD3-dependent sites of m^2^G within polyA+ enriched RNA from A549 lung cancer cells, which contain predominantly mRNAs and long non-coding RNAs. Our detailed analysis revealed a list, mainly of mRNAs, containing enzyme-dependent sites of m^2^G. Many of these transcripts are associated with human diseases, in particular cancer, which is of particular note due to the association of THUMPD3 with human cancers. We anticipate that these data will provide a valuable resource for other researchers in the field and support seminal studies in the role of m^2^G in human health and disease. In addition, our approach identified many other putative m^2^G sites within the transcriptome, which appear not to be installed by THUMPD3. These sites represent a useful database for workers in the RNA modification field, particularly those analysing THUMPD2 and TRMT11. It is likely that our initial list of m^2^G-modified sites within transcripts represents a significant underestimate of the true number. This is largely because of inherent limitations in our ability to differentiate signal-to-noise in our analyses. To increase this ratio, future approaches could include depletion of other m^2^G methyltransferases, either singularly or in combination.

The bioinformatic pipeline we have developed to detect THUMPD3-dependent m^2^G sites can be readily adapted to detect other photooxidation induced mutations at modified G. For instance, the pipeline can be applied to detect other methylated modifications, such as m^2^_2_G or m^1^G. Indeed, PhOxi-seq may be feasible to determine m^1^G sites using known demethylase enzymes; this approach is currently being investigated and will be reported in a forthcoming study. Lastly, the pipeline developed here can also be easily tailored to detect single-nucleotide changes induced by any method at any nucleotide, making it highly versatile.

In summary, our application of PhOxi-seq has unveiled numerous potential sites of m^2^G, including THUMPD3-dependent and independent targets, across mRNA regions and non-coding RNAs. These findings emphasize the diverse landscape of m^2^G within the transcriptome. Additionally, the data will support further exploration into the functional implications of these modified sites in RNA biology.

## ASSOCIATED CONTENT

### Supporting Information

The Supporting Information is available free of charge on the ACS Publications website.

Experimental methods; sequencing and data analysis; supporting data (Figures S1-S5) and sequencing data (Tables S1-S4)

## AUTHOR INFORMATION

### Notes

Any additional relevant notes should be placed here.

## ACKNOWLEDGMENT

The T.K. laboratory is supported by grants from Cancer Research UK (RG96894 and C6946/A24843) and the Wellcome Trust (WT203144). M.K. received PhD studentship funding from CRUK (RG96894) and Storm Therapeutics. The R.H. laboratory is supported by the Natural Sciences and Engineering Research Council of Canada, Ontario Ministry of Research and Innovation, Canada Foundation for Innovation, and York University.

---

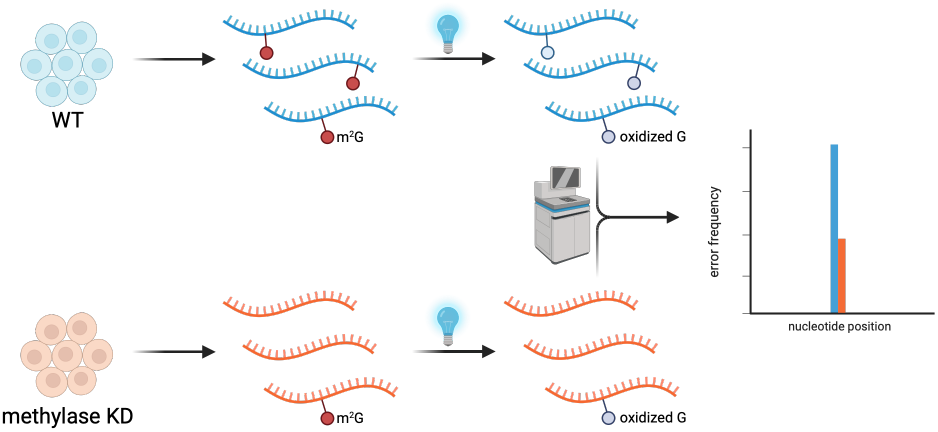

## REFERENCES

(1) Boccaletto, P.; Stefaniak, F.; Ray, A.; Cappannini, A.; Mukherjee, S.; Purta, E.; Kurkowska, M.; Shirvanizadeh, N.; Destefanis, E.; Groza P.; Avşar, G.; Romitelli, A.; Pir, P.; Dassi, E.; Conticello, S. G.; Aguilo, F.; Bujnicki, J. M. MODOMICS: a database of RNA modification pathways. 2021 update. Nucleic Acids Res. 2022, 50 (D1), D231–D235.

(2) Zhao, B. S.; Roundtree, I. A; He, C. Post-transcriptional gene regulation by mRNA modifications. Nat. Rev. Mol. Cell Biol. 2017, 18, 31–42.

(3) Barbieri, I.; Kouzarides, T. Role of RNA modifications in cancer. Nat Rev Cancer 2020, 20 (6), 303–322.

(4) Bartee, D.; Gamage, S. T.; Link, C. N.; Meier, J. L. Arrow pushing in RNA modification sequencing. Chem. Soc. Rev. 2021, 50, 9482–9502.

(5) Li, X.; Xiong, X.; Yi, C. Epitranscriptome sequencing technologies: decoding RNA modifications. Nat. Meth. 2017, 14, 23–31.

(6) Jiang, X.; Liu, B.; Nie, Z.; Duan, L.; Xiong, Q.; Jin, Z.; Yang, C.; Chen, Y. The role of m6A modification in the biological functions and diseases. Signal Transduct. Target. Ther. 2021, 6, 74.

(7) Jones, J. D.; Franco, M.; Smith, T.; Snyder, L. R.; Anders, A. G.; Ruotolo, B. T.; Kennedy, R. T.; Koutmou, K. S. Methylated guanosine and uridine modifications in S. cerevisiae mRNAs modulate translation elongation. RSC Chem. Biol. 2023, 4, 363–378.

(8) Yang, W.-Q.; Xiong, Q.-P.; Ge, J.-Y.; Li, H.; Zhu, W.-Y.; Nie, Y.; Lin, X.; Lv, D.; Li, J.; Lin, H.; Liu, R.-J. THUMPD3-TRMT112 is a m2G methyltransferase working on a broad range of tRNA substrates. Nucleic Acids Res. 2021, 49 (20), 11900–11919.

(9) Wang, C.; Ulryck, N.; Herzel, L.; Pythoud, N.; Kleiber, N.; Guérineau, V.; Jactel, V.; Moritz, C.; Bohnsack, M. T.; Carapito, C.; Touboul, D.; Bohnsack, K. E.; Graille, M. N2-methylguanosine modifications on human tRNAs and snRNA U6 are important for cell proliferation, protein translation and pre-mRNA splicing. Nucleic Acids Res. 2023, 51 (14), 7496–7519.

(10) Yang, W.-Q.; Ge, J.-Y.; Zhang, X.; Zhu, W.-Y.; Lin, L.; Shi, Y.; Xu, B.; Liu, R.-J. THUMPD2 catalyzes the N2-methylation of U6 snRNA of the spliceosome catalytic center and regulates pre-mRNA splicing and retinal degeneration. Nucleic Acids Res. 2024, 52 (6), 3291–3309.

(11) Klimontova, M.; Zhang, H.; Campos-Laborie, F.; Webster, N.; Andrews, B.; Chung Kim Chung, K.; Hili, R.; Kouzarides, K.; Bannister, A. J. THUMPD3 regulates alternative splicing of ECM transcripts in human lung cancer cells and promotes proliferation and migration.

(12) Yu, Y.-P.; Ding, Y.; Chen, Z.; et al. Novel fusion transcripts associate with progressive prostate cancer. Am. J. Pathol. 2014, 184, 2840–2849.

(13) Zheng, H.-X.; Zhang, X.-S.; Sui, N. Advances in the profiling of N6-methyladenosine (m6A) modifications. Biotechnol. Adv. 2020, 45, 107656.

(14) Chung Kim Chung, K.; Mahdavi-Amiri, Y.; Korfmann, C.; Hili, R. PhOxi-Seq: Single-nucleotide resolution sequencing of N2-methylation at guanosine in RNA by photoredox catalysis. J. Am. Chem. Soc. 2022, 144, 5723–5727.

(15) Batista, P. J.; Molinie, B.; Wang, J.; Qu, K.; Zhang, J.; Li, L.; Bouley, D. M.; Lujan, E.; Haddad, B.; Daneshvar, K. et al. m6A RNA Modification Controls Cell Fate Transition in Mammalian Embryonic Stem Cells. Cell Stem Cell 2014, 15, 707–719.

(16) Schwartz, S.; Mumbach, M. R.; Jovanovic, M.; Wang, T.; Maciag, K.; Bushkin, G. G. et al. Perturbation of m6A writers reveals two distinct classes of mRNA methylation at internal and 5’ sites. Cell Rep. 2014, 8, 284–296.

(17) Geula, S.; Moshitch-Moshkovitz, S.; Dominissini, D.; Mansour, A. A.; Kol, N. Salmon-Divon M, et al. m6A mRNA methylation facilitates resolution of naïve pluripotency toward differentiation. Science. 2015, 347, 1002–1006.

(18) Su, R.; Dong, L.; Li, Y.; Gao, M.; He, P. C.; Liu, W.; et al. METTL16 exerts an m6A-independent function to facilitate translation and tumorigenesis. Nat Cell Biol. 2022, 24, 205–216.

(19) Wei, G.; Almeida, M.; Pintacuda, G.; Coker, H.; Bowness, J. S.; Ule, J.; et al. Acute depletion of METTL3 implicates N6-methyladenosine in alternative intron/exon inclusion in the nascent transcriptome. Genome Res. 2021, 31, 1395–1408.

(20) Shigematsu, M.; Honda, S.; Loher, P.; Telonis, A. G.; Rigoutsos, I.; Kirino, Y. YAMAT-seq: an efficient method for high-throughput sequencing of mature transfer RNAs. Nucleic Acids Res. 2017, 45, No. e70.

(21) Poh, H. X.; Mirza, A. H.; Pickering, B. F.; Jaffrey, S. R. Alternative splicing of METTL3 explains apparently METTL3-independent m6A modifications in mRNA. Plos Biol. 2022, 20, No. e3001683.

(22) Huang, W.; Fan, L.; Tang, Y.; Chi, Y.; Li, J. A pan-cancer analysis of the oncogenic role of integrin beta4 (ITGB4) in human tumors. Int. J. Gen. Med. 2021, 14, 9629–9645.

(23) Li, G.; Yi, S.; Yang, F.; Zhou, Y.; Ji, Q.; Cai, J.; Mei, Y. Identification of mutant genes with high-frequency, high-risk, and high-expression in lung adenocarcinoma. Thorac. Cancer 2014, 5, 211–218.

(24) Lee, K. K.; Rajagopalan, D.; Bhatia, S. S.; Tirado-Magallanes, R.; Chng, W. J.; Jha, S. The oncogenic E3 ligase TRIP12 suppresses epithelial–mesenchymal transition (EMT) and mesenchymal traits through ZEB1/2. Cell Death Disc. 2021, 7, 95.

