## Supporting Information for "PhOxi-seq detects enzyme-dependent m^2^G in multiple RNA types"

1. **Cell culture, RNA purification and sequencing**

### 1.1. Cell culture media, reagents and maintenance

All cell culture was carried out under sterile conditions in a standard laminar flow hood. Human cell lines were maintained in filter top flasks or culture dishes and cultured in DMEM (Gibco), 10% [v/v] FBS growth media. 293T and A549 cell lines were grown in DMEM supplemented with 1x penicillin-streptomycin (pen-strep) antibiotics and 10% [v/v] fetal bovine serum (FBS). Cells were grown at 37˚C, 5% [v/v] $CO_2$ and were maintained at > 80% viability. Cells were passaged every 3-5 days and seeded at 1 x $10^5$ live cells per ml. Cells were passaged using conventional cell culture techniques. The maximum number of passages was 20 before a new vial of cells was revived. Tests for mycoplasma contamination were carried out every month or when a new vial of cells was revived.

### 1.2. Lentiviral production and infection

The LentiCRISPR v2 system was used to generate CRISPR/Cas9 knockout cell lines of RNA methyltransferases THUMPD3.[1] The 'all-in-one' pLKO-Tet-On system was used to generate inducible shRNA-mediated THUMPD3 knockdowns.[2,3] Constructs carrying sgRNAs and shRNAs are listed in Table 1. 293T cells were seeded onto L-polylysine (1 µg/ml) coated 10 cm dishes 24 hours prior to transfection in antibiotic-free DMEM media with 10% [v/v] FCS. Cells were 80% confluent on the day of transfection. To produce lentiviral particles, cells were transfected using 42 µl FuGENE® 4K Transfection Reagent (Promega, E5911) with 6 µg of lentiCRISPR v2 or pLKO-Tet-On constructs together with 3.5 µg psPAX2 (Addgene, 12260) and 4 µg pCMV-VSV-G (Addgene, 8454). 16 hours later, the media was exchanged for fresh media. 48 hours post-infection, virus particles were harvested and sterile filtered using 0.45 µm syringe filters (Millipore). Aliquots of lentiviral supernatants were stored at -70˚C.

Target cells (A549) were then transduced with the virus with polybrene (8 µg/ml) to increase the efficiency of transduction. 48 hours after infection transduced cells were selected for by treatment with puromycin (1 µg/ml). To induce expression of shRNAs, 10-100 ng/ml doxycycline was added to the media and the cells were incubated for 4-8 days to allow adequate expression of the shRNA.

### 1.3. Total RNA extraction with DNase treatment

Approximately $10^6$ cells were homogenized in 1 ml QIAzol lysis reagent (QIAGEN). Homogenates were incubated for 5 minutes at room temperature. RNA was supplemented with chloroform (0.2 ml per 1 ml of QIAzol) and vigorously shaken for 20 seconds. After 3 minutes incubation at room temperature, homogenates were centrifuged at 12000 g for 15 minutes, at 4˚C. The aqueous (top) phase was carefully removed and transferred to a new RNase-free tube. 1.5 volumes of 100% EtOH were added. RNA was purified using a RNeasy Mini Kit (QIAGEN, 74104) with an additional step of DNase digestion after the first wash following the RNase-Free

DNase Set protocol (QIAGEN, 79254). After the washes, columns were centrifuged for an additional 2 minutes to remove residual washing buffer. RNA was eluted with 30-50 µl of RNase-free water. RNA concentrations were measured using a Nanodrop spectrophotometer (Thermo Fisher) or Qubit BR RNA (Thermo Fisher, Q10211). RNA was stored at -70˚C until further use.

Table 1. Plasmids used to generate CRISPR KO and shRNA-inducible knockdown cells.

| Name | Vector | Insert |
|---|---|---|
| sgscr | lentiCRISPR v2 | **fwd** - CACCGGCGAGGTATTCGGCTCCGCG<br>**rev** - AAACCGCGGAGCCGAATACCTCGCC |
| sgD3 #3 | lentiCRISPR v2 | **fwd** - CACCGCGCAAGAGGCCAATGGAGTG<br>**rev** - AAACCACTCCATTGGCCTCTTGCGC |
| shscr | pLKO-Tet-On | **fwd** - CCGGCCTAAGGTTAAGTCGCCCTCGCTC GAGCGAGGGCGACTTAACCTTAGGTTTTT<br>**rev** - AATTAAAAACCTAAGGTTAAGTCGCCCTCG CTCGAGCGAGGGCGACTTAACCTTAGG |
| shD3 #1 | pLKO-Tet-On | **fwd** - CCGGCCTGAAGGAATGAAGCATTAT CTCGAGATAATGCTTCATTCCTTCAGGTTTTT<br>**rev** - AATTAAAAACCTGAAGGAATGAAGCATTAT CTCGAGATAATGCTTCATTCCTTCAGG |

### 1.4. Purification of small and large RNA fractions

RNA Clean & Concentrator kits (ZYMO RESEARCH, R1013 or R1017) were used to separate small (< 200nt) and large (> 200nt) RNA fractions, following the manufacturer's instructions. RNA concentration was determined using a QubitTM RNA HS Assay Kit (Thermo Fisher, Q32855). RNA was stored at -70˚C until needed.

### 1.5. PhOxi-seq on tRNA

A549 CRISPR/Cas9 targeted KO cell lines (sgscr and sgD3 #3, Table 1) were grown to 70-80% confluency. Cells were then harvested by scraping and pellets. Total RNA was then extracted from the cells as described above (Section 1.3). Next, the small RNA fraction was purified (above, Section 1.4) and subjected to blue light induced photo-oxidation (Section 1.7). Primers that specifically amplify tRNA-Gly-GCC-2 and tRNA-Gly-CCC-2 were used prior to YAMAT-seq.[4] Three biological replicates were included for each treatment. Bioinformatic analysis was performed as described in Section 2.2.

### 1.6. PhOxi-seq using mRNA as a template

A549 cell lines harboring shRNA vectors (shscr and shD3 #1, Table 1) were seeded into 15 cm dishes. shRNA expression was induced by doxycycline for 8 days. Cell pellets were collected, and total RNA was extracted as described above. Photo-oxidation was performed as described in Section 1.7. rRNA HMR Removal Kit (Qiagen, 334386) was used in conjunction with

NEXTFLEX® Rapid Directional RNA-Seq Kit 2.0 (PerkinElmer, NOVA-5198-02) to deplete rRNA and perform whole transcriptome analysis according to the manufacturer's instructions. The RNA quality was assessed using an RNA Screen Tape (Agilent, 5067-5576) analyzed on a 4200 TapeStation System (Agilent, G2991BA). The NEXTFLEX® Rapid Directional RNA-Seq Kit 2.0 protocol was followed to perform all the remaining steps for library construction steps. The final multiplexed library, with an average size of 350-550 bp (at 10-20 nM) was submitted for 150 bp, paired-end sequencing on an Illumina NovaSeq® instrument. To ensure deep sequencing, a maximum of six samples was sequenced at each time. 120 GB of raw data were obtained per multiplexed library. Bioinformatic analyses was performed as describe in Section 2.

#### 1.7. PhOxi-seq with RNA

The photocatalysis reaction was performed in a PCR tube by adding 5 $\mu$L of 2 $\mu$M RNA [in nuclease-free water (Ambion, AM9937)], 13.2 $\mu$L of 1 mM selectfluor (in 100 mM phosphate buffer, pH 7), 1.2 $\mu$L of 0.5 mM riboflavin (in 100 mM phosphate buffer, pH 7), and phosphate buffer (100 mM, pH 7) up to 30 $\mu$L. Reaction was left at 65 cm from blue light (Kessil Tune Blue A160WE lamp set to lowest intensity) for 2 h, followed by purification with Monarch RNA cleanup kit.

### 2. Data analysis

#### 2.1. tRNA sequencing analysis

##### 2.1.1. Data processing

Read quality was assessed using FastQC (v0.11.9).[5] Reads were then filtered to include reads between 103 and 123 base pair (bp) long (113± 10 bp) using custom bash (v5.0.17) scripts. Then reads were filtered to include mature tRNA reads that end in 'GGA' or CCA' using custom bash scripts.

Filtered reads were then aligned to the tRNA reference sequence using bwa-mem (v0.7.17)[6] with parameters -t 32 -w 20. Then, aligned reads were filtered to include only primarily mapped reads using samtools (v1.10)[7] view with parameter -F 260. Hard-clipped reads were also removed using samtools. The filtered reads were then sorted and indexed using samtools. The proportion of different bases or indels at each nucleotide position of the tRNA sequence for aligned reads of each tRNA isodecoder type were summarised using bam-readcount (v1.0.1).[8]

##### 2.1.2. tRNA reference sequence

A comprehensive and non-repetitive set of reference sequences of tRNAs was constructed. It consists of isodecoder sequences of high confidence mature nuclear-encoded cytoplasmic tRNA genes (n=432) from GtRNAdb[9] and mitochondrial tRNA genes (n=28) from tRNAdb.[10]

### 2.2. PhOxi-seq pipeline based on transcriptome-wide analysis

#### 2.2.1. Data processing

Adapter sequences were trimmed from 150bp paired-end reads using Trimmomatic (v0.40-rc1).[11] Filtered reads were aligned using STAR (v2.7.10a)[12] two-pass mode with alignment parameters --outFilterMultimapNmax 20 --alignSJoverhangMin 5 --alignSJDBoverhangMin 3 --outFilterMismatchNmax 999 --outFilterMismatchNoverReadLmax 1.0 --alignIntronMin 21 --sjdbOverhang 149 --alignIntronMax 1000000 --alignMatesGapMax 1000000. Aligned reads were then filtered to include only primarily mapped reads using samtools (v1.10) view with parameter -F 260. Biological replicates (n=3) for treated and untreated samples were merged respectively using samtools merge. Reads spanning introns was reformatted using SplitNCigarReads from GATK (v4.3.0.0).[13] The variant calling was performed for merged reads using bcftools (v1.16) mpileup with parameters --max-depth 1000000 and bcftools call under multiallelic mode[14] with parameters –multiallelic-caller –variants-only. The resulting vcf file containing variant information was processed using bedtools (v2.30.0)[15] intersect program to obtain bed file of genomic regions with variant calls. bam-readcount (v1.0.1). The subsequent analysis and plotting were performed using custom Python (v3.8) scripts.

#### 2.2.2. Data exploration

The overall distribution of variant allele frequency (VAF) data of variant sites from bcftools (with non-zero depth in all replicates) (n=1192693) was investigated. Density plots of the overall distribution for every replicate from each experiment group were plotted to observe if BL treatment and enzyme depletion influenced the overall distribution of the data.

In addition, Pearson's correlation coefficient between VAF data between all samples was calculated and plotted as heatmap. It shows a global similarity of overall data between samples.

#### 2.2.3. Filtering conditions for potential $m^2G$ sites

The filtering conditions to select top potential m2G modified sites focused on these aspects:

1. Sufficient sequencing coverage
2. Multiallelic or deletion signature
3. Levels of sequencing errors increase upon BL treatment
4. Levels of sequencing errors decrease upon enzyme depletion

The conditions are based on the observation of positive samples in tRNA sequence analysis. The cutoff of levels of sequencing errors it follows the assumptions that the levels of sequencing errors in control samples should be low since m2G is non base pair-disrupting modification. Also, the cutoff of sequencing error rate is non-stringent allowing more potential sites to be included.

To define the cutoff of sequencing coverage, we calculated correlation between different samples upon different sequencing coverage. The average sequencing depth of all 'SNP' sites across different samples were calculated. These sites are then sorted, in ascending order, by coverage. A non-overlapping sliding window approach is used to group the sorted sites where sites with similar coverage in a specific range are grouped into 'windows'. For each window (e.g. log(depth): 2-3, 3-4), the correlation of the VAF data of the sites from the sample of interest and all other samples is calculated and averaged. Per sample, the correlation with the rest of samples (y-axis) is represented versus the average window coverage (x-axis). Around the depth of 50, the values of correlation coefficients plateau.

#### 2.3. Analysis of sequencing error rates of target sites

For selected top targets of the whole transcriptome, sequencing error data of genomic region 50bp upstream to 50bp downstream of the target site were extracted and guanosine errors were plotted. In terms of tRNAs, sequencing error data of complete sequence positions of tRNAs were extracted and guanosine errors were plotted. Specifically, sequencing error changes upon BL treatment in control samples and sequencing error changes upon enzyme depletion in BL treated samples were plotted in bar plots.

In tRNA sequencing analysis, levels of main error types (G->T substitution, G->C substitution, G->A substitution and single base deletion) with and without BL treatment in control samples were plotted respectively for two tRNA isodecoder types (tRNA-Gly-GCC-2 and tRNA-Gly-CCC-2).

Replicate information was also included when analysing sequencing error levels at nucleotide level. In this case, reads from individual replicate were processed through the pipeline instead of merging the replicates for deeper sequencing coverage and used for downstream analysis.

#### 2.4. Statistical analysis

Welch's t-test was used to compare sequencing error levels (n=3) between different samples given that the data follows the normality assumption but with different variances. Pearson's correlation coefficients were calculated to compare the similarity of overall VAF data between samples. Spearman's rank correlation coefficient was calculated for similarity between data of different samples in different depth ranges.

#### 2.5. Code availability

The computational methods and custom scripts are available in the following Github repository: https://github.com/hanzhang2000/PhOxi_seq_transcriptome_code.

### 3. Supporting data

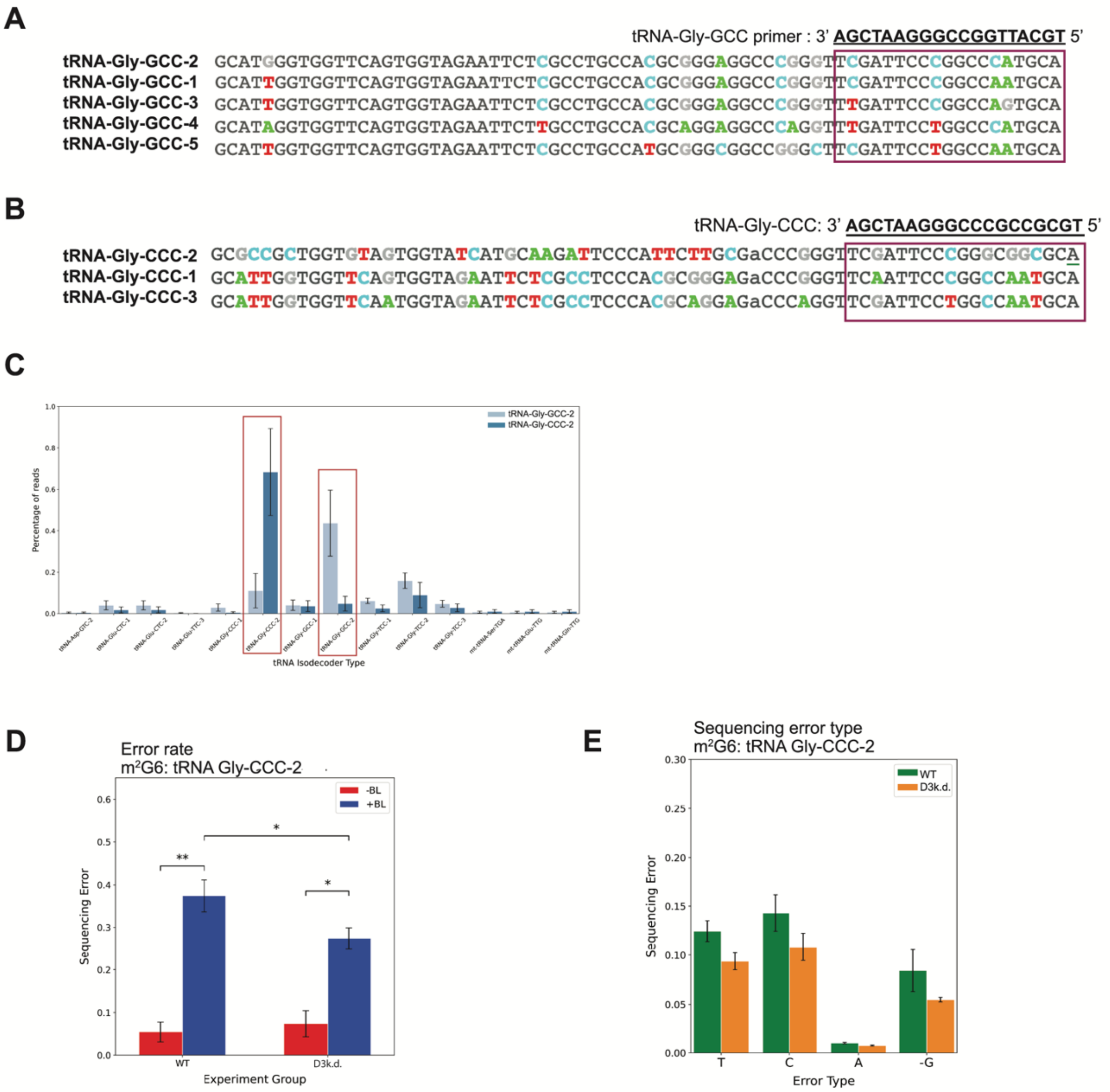

**Figure S1. tRNA sequencing, mapping, and sequencing profile of another tRNA type. (A)** Annealing of unique tRNA-Gly-GCC-2 RT-PCR primer to five tRNA-Gly-GCC isodecoders. **(B)** Annealing of unique tRNA-Gly-CCC-2 RT-PCR primer to three tRNA-Gly-CCC isodecoders. **(C)** Reads acquired from YAMAT-seq were aligned to a comprehensive set of nuclear-encoded cytoplasmic tRNA and mitochondrial tRNA sequences. The top tRNAs that the reads were predominantly aligned to are depicted. The data is illustrated as the proportion of reads, where 0 represents 0% and 1.0 signifies 100%. Error bars represent the mean ± SD of 3 independent replicates (n=3). **(D)** Plot of guanosine error rates for tRNA-Gly-CCC-2 in WT cells. Red bars indicate sequencing run without photo-oxidation (-BL) and blue bars indicate PhOxi-seq run (+BL). The letters on the x-axis correspond to the nucleotides of tRNA, beginning with the first nucleotide. **(E)** Plot of guanosine errors for the photo-oxidised tRNA-Gly-CCC-2. Green bars indicate WT samples and orange bars indicate D3k.d. samples. The sequencing error is illustrated in proportion, where 0 corresponds to 0% and 1 represents 100%. **(F)** Plot representing sequencing error at the position G6 with (+BL) and without photo-oxidation (-BL). Two experimental groups represent WT and D3k.d. samples. Welch's t-test conducted to assess the significance of changes in error rates at G6 on tRNA-Gly-CCC-2. **(G)** Plot representing types of sequencing error occurring at the G6 position on tRNA-Gly-CCC-2 after photo-oxidation. Green bars denote WT cells, orange bars denote D3k.d. cells.

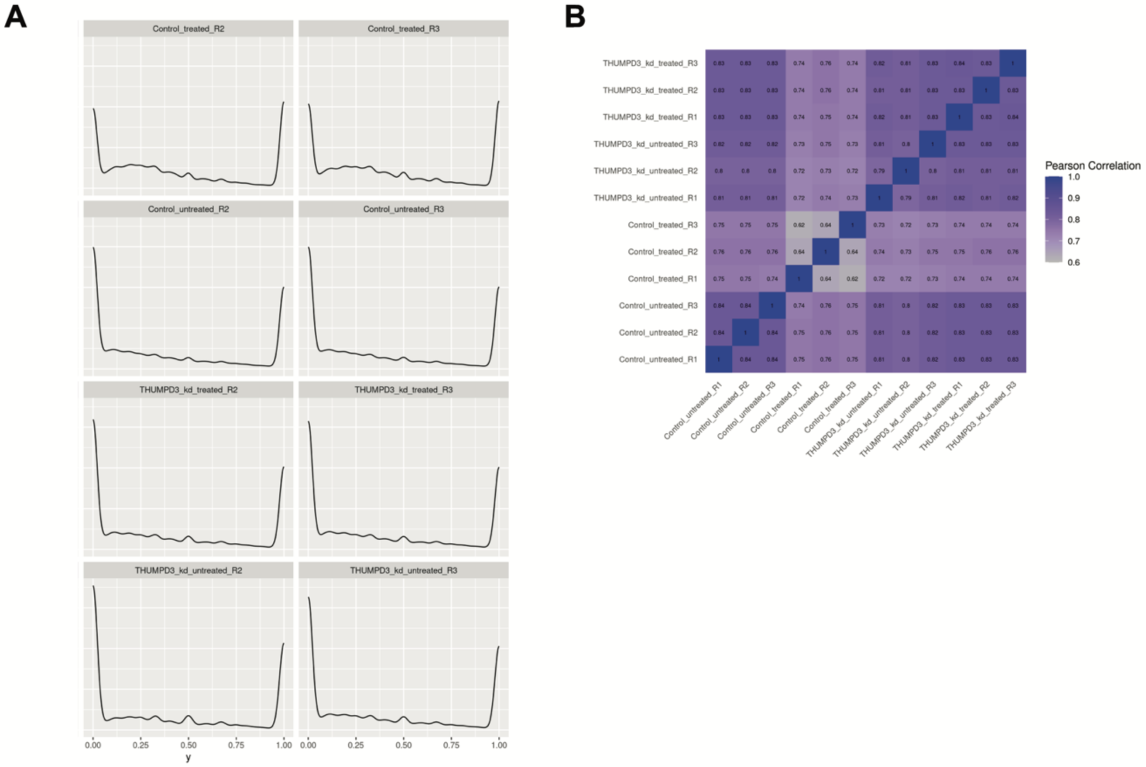

**Figure S2. Additional QC measures. (A)** Remaining density plots (additional two biological replicates) illustrating overall distribution of variant allele frequency (VAF) data of variant sites of representative replicate from each experiment group (control cell line (WT) with and without BL (±BL) treatment, THUMPD3 depleted cell line (D3k.d.) ±BL treatment). The y-axis of the density curve is density and the area under the curve gives the percentage. The x-axis ranges from 0-1, representing the VAF data, which here indicates the proportion of reads that do not match the reference genome at each variant site. The consistent pattern in the plots with two peaks correspond to specific sets of data: one peak with VAF values (x-axis) approaching 1 represent mostly SNPs; one with VAF values approaching 0 could be mostly noises. The VAF data in this plot covers variant sites from the variant calling step that show non-zero depth in all replicates (n=1192693). **(B)** Heatmap based on Pearson Correlation coefficient of VAF data between samples. The VAF data here covers variant sites that show non-zero depth in all replicates (n=1192693).

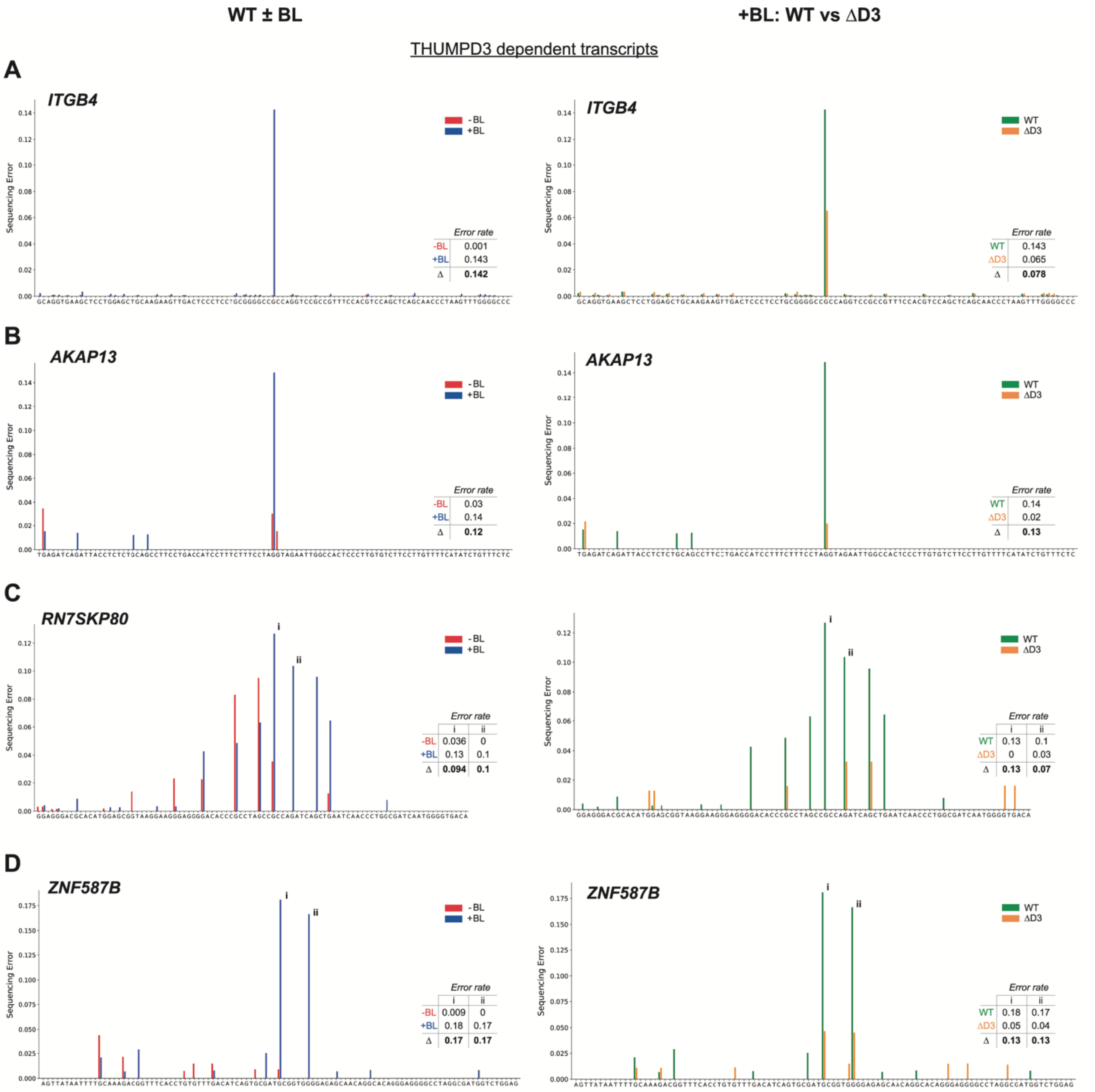

**Figure S3. Effect of PhOxi-seq treatment on total sequencing errors in the genomic regions of top potential THUMPD3 dependent m$^2$G sites.** Bar plots showing total sequencing errors/VAF (y-axis) in the region 50bp upstream to 50bp downstream (x-axis) of potential m$^2$G site. **Left panel:** bar plots represent guanosine error rates in WT cells with (+BL, depicted in blue) and without (-BL, depicted in red). **Right panel:** plot of guanosine errors for the photo-oxidised samples (+BL). Green bars indicate WT cell lines and orange bars indicate THUMPD3 depleted cell line (D3k.d.). The sequencing error is illustrated in percentage, where 0 is 0% and 1 represents 100%. **Insert in graph:** tables indicating error rates at given m$^2$G sites. <u>THUMPD3-dependent targets:</u> **(A)** exon of *ITGB4*; **(B)** intron of *AKAP13*; **(C)** *RN7SKP80*, pseudogene of snRNA *7SK*; **(D)** 3' UTR of *ZNF587F*.

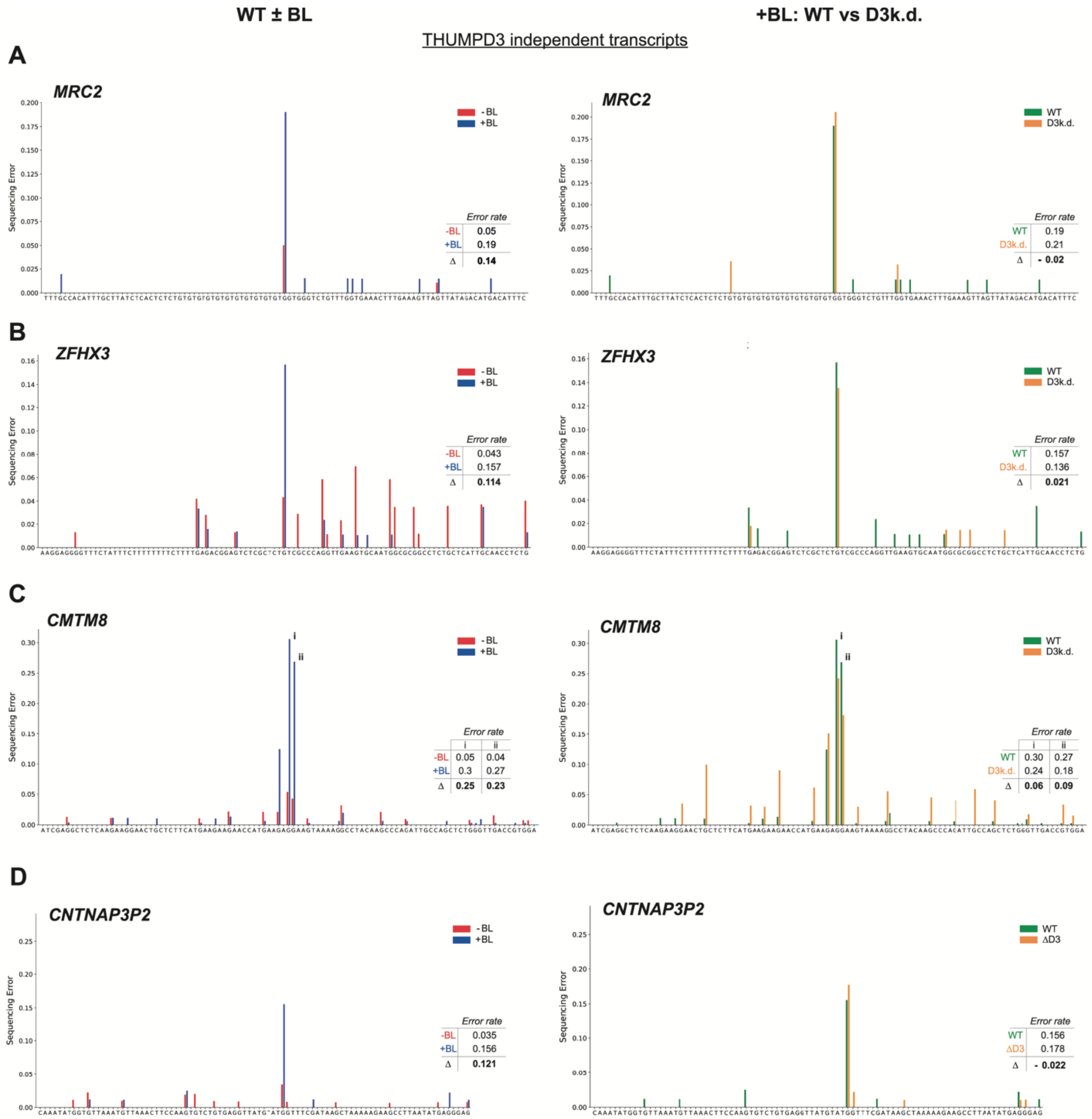

**Figure S4. Effect of PhOxi-seq treatment on total sequencing errors in the genomic regions of alternative sets of top potential m$^2$G sites that do not exhibit THUMPD3 dependency.** Bar plots showing total sequencing errors/VAF (y-axis) in the region 50bp upstream to 50bp downstream (x-axis) of potential m$^2$G site. **Left panel:** bar plots represent guanosine error rates in WT cells with (+BL) and without (-BL). **Right panel:** plot of guanosine errors for the photo-oxidised samples (+BL). Green bars indicate WT samples and orange bars indicate D3k.d. samples. The sequencing error is illustrated in percentage, where 0 is 0% and 1 represents 100%. **Insert in graph:** tables indicating error rates at given m$^2$G sites. **(A)** Intron of *MRC2*; **(B)** intron of *ZFHX3*; **(C)** intron of *CMTM8*. **(D)** *CNTNAP3P2*.

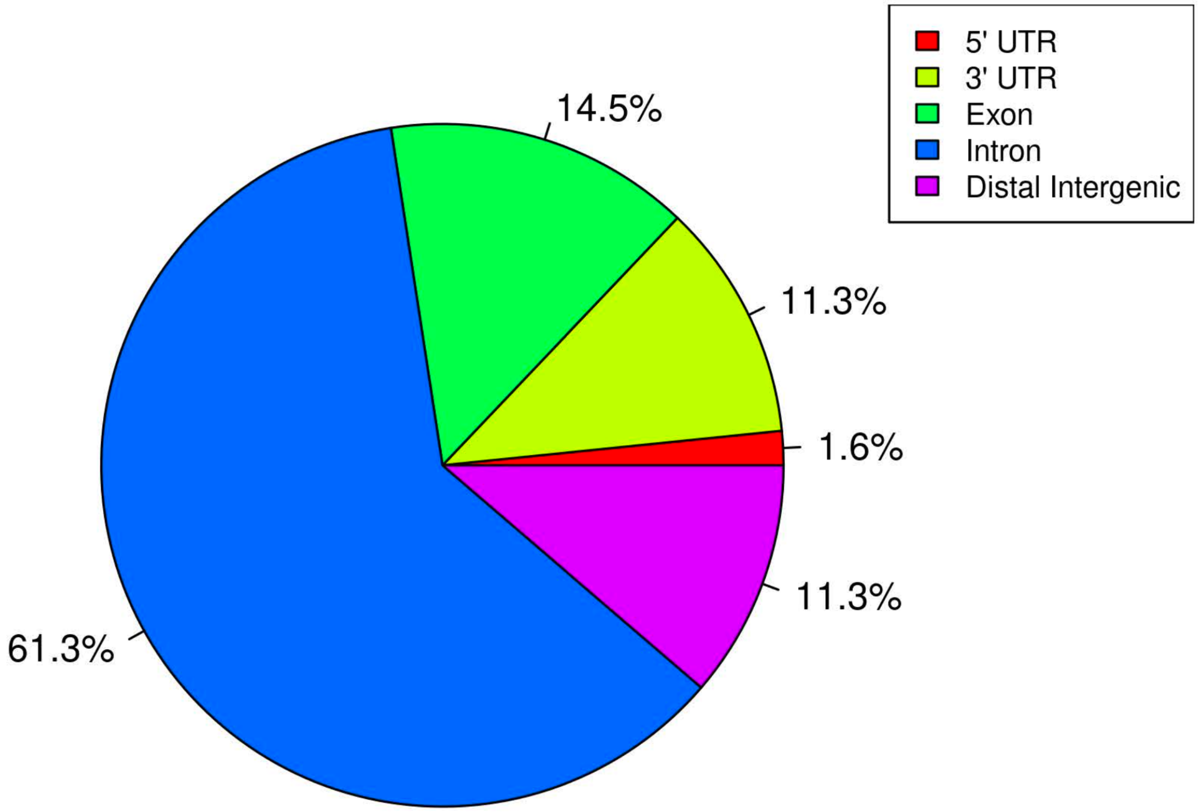

**Figure S5. Distribution of putative m$^2$G sites in mRNA**

### 4. References

[1] Shalem, O.; Sanjana, N. E.; Hartenian, E.; Shi, X.; Scott, D. A.; Mikkelson, T.; Heckl, D.; Ebert, B. L.; Root, D. E.; Doench, J. G.; Zhang, F. Genome-scale crispr-cas9 knockout screening in human cells. *Science* **2014**, *343* (6166), 84–87.
[2] Wiederschain, D.; Wee, S.; Chen, L.; Loo, A.; Yang, G.; Huang, A.; Chen, Y.; Caponigro, G.; Yao, Y.-M.; Lengauer, C.; Sellers, W. R.; Benson, J. D. Single-vector inducible lentiviral rnai system for oncology target validation. *Cell Cycle* **2009**, *8* (3), 498–504.
[3] Wee, S.; Wiederschain, D.; Maira, S.-M.; Loo, A.; Miller, C.; deBeaumont, R.; Stegmeier, F.; Yao, Y.-M.; Lengauer, C. PTEN-deficient cancers depend on PIK3CB. *Proc. Natl. Acad. Sci. U.S.A.* **2008**, *105* (35), 13057–13062.
[4] Shigematsu, M.; Honda, S.; Loher, P.; Telonis, A. G.; Rigoutsos, I.; Kirino, Y. Yamat- seq: an efficient method for high-throughput sequencing of mature transfer RNAs. *Nucleic Acids Res.* **2017**, *45* (9), No. e70.
[5] Andrews S. (2010). FastQC: a quality control tool for high throughput sequence data. Available online at http://www.bioinformatics.babraham.ac.uk/projects/fastqc
[6] Li H. Aligning sequence reads, clone sequences and assembly contigs with BWA-MEM. 2013, arXiv:1303.3997v2[q-bio.GN]. arXiv.org e-Print archive. https://doi.org/10.48550/arXiv.1303.3997
[7] Danecek, P.; Bonfield, J. K.; Liddle, J.; Marshall, J.; Ohan, V.; Pollard, M. O.; Whitwham, A.; Keane, T.; McCarthy, S. A.; Davies, R. M.; Li, H. Twelve years of SAMtools and BCFtools, *GigaScience* **2021**, *10* (2), giab008.
[8] Khanna, A.; Larson, D. E.; Srivatsan, S. N.; Mosior, M.; Abbott, T. E.; Kiwala, S.; et al. Bam-readcount - rapid generation of basepair-resolution sequence metrics. *J. Open Source Softw*. **2022**, *7* (69), 3722.
[9] Chan, P. P.; Lowe, T. M. GtRNAdb 2.0: an expanded database of transfer RNA genes identified in complete and draft genomes. *Nucleic Acids Res*. **2016**, *44* (D1), D184–189.
[10] Jühling, F.; Mörl, M.; Hartmann, R. K.; Sprinzl, M.; Stadler, P. F.; Pütz, J. tRNAdb 2009: compilation of tRNA sequences and tRNA genes. *Nucleic Acids Res*. **2009**, *37*, D159–162.
[11] Bolger, A. M.; Lohse, M.; Usadel, B. Trimmomatic: a flexible trimmer for Illumina sequence data. *Bioinformatics* **2014**, *30* (15), 2114–20.
[12] Dobin, A.; Davis, C. A.; Schlesinger, F.; Drenkow, J.; Zaleski, C.; Jha, S.; et al. STAR: ultrafast universal RNA-seq aligner. *Bioinformatics* **2013**, *29* (1), 15–21.
[13] McKenna, A.; Hanna, M.; Banks, E.; Sivachenko, A.; Cibulskis, K.; Kernytsky, A.; et al. The Genome Analysis Toolkit: A MapReduce framework for analyzing next-generation DNA sequencing data. *Genome Res*. **2010**, *20* (9), 1297–1303.
[14] Danecek, P.; Schiffels, S.; Durbin, R. Multiallelic calling model in bcftools (-m). https://samtools.github.io/bcftools/call-m.pdf
[15] Quinlan, A. R.; Hall, I. M. BEDTools: a flexible suite of utilities for comparing genomic features. *Bioinformatics* **2010**, *26* (6), 841–842.
